# Modeling aging and retinal degeneration with mitochondrial DNA mutation burden

**DOI:** 10.1101/2023.11.30.569464

**Authors:** John Sturgis, Rupesh Singh, Quinn Caron, Ivy S. Samuels, Thomas Micheal Shiju, Aditi Mukkara, Paul Freedman, Vera L. Bonilha

## Abstract

Somatic mitochondrial DNA (mtDNA) mutation accumulation has been observed in individuals with retinal degenerative disorders. To study the effects of aging and mtDNA mutation accumulation in the retina, a Polymerase gamma (POLG) deficiency model, the POLG^D257A^ mutator mice (PolgD257A), was used. POLG is an enzyme responsible for regulating mtDNA replication and repair. Retinas of young and older mice with this mutation were analyzed in vivo and ex vivo to provide new insights into the contribution of age-related mitochondrial dysfunction due to mtDNA damage. Optical coherence tomography (OCT) image analysis revealed a decrease in retinal and photoreceptor thickness starting at 6 months of age in mice with the POLG^D257A^ mutation compared to wild-type (WT) mice. Electroretinography (ERG) testing showed a significant decrease in all recorded responses at 6 months of age. Sections labeled with markers of different types of retinal cells, including cones, rods, and bipolar cells, exhibited decreased labeling starting at 6 months. However, electron microscopy analysis revealed differences in retinal pigment epithelium (RPE) mitochondria morphology beginning at 3 months. Interestingly, there was no increase in oxidative stress observed in the retina or RPE of POLGD257A mice. Additionally, POLGD257A RPE exhibited an accelerated rate of autofluorescence cytoplasmic granule formation and accumulation. Mitochondrial markers displayed decreased abundance in protein lysates obtained from retina and RPE samples. These findings suggest that the accumulation of mitochondrial DNA mutations leads to impaired mitochondrial function and accelerated aging, resulting in retinal degeneration.

## 1. INTRODUCTION

Mitochondria are unique eukaryotic organelles because they possess their own mitochondrial genome, separate from the nuclear genome. The mitochondrial genome contains approximately 16.5 kilobases; it encodes 37 genes, including 13 proteins, 22 transfer RNAs (tRNA), and two ribosomal RNAs (rRNA). These genes encode the subunits of the electron transport chain (ETC), which is responsible for oxidative phosphorylation and energy production (Chinnery & Hudson, 2013; Pinto & Moraes, 2015). Mitochondrial genetic instability can occur through various pathways, such as base-substitution mutations, deletions, failure of base-excision repair and oxidative insult. (Kujoth et al., 2007). Additionally, mitochondrial DNA (mtDNA) is more susceptible to damage due to its high replication rate, proximity to free radical production sites, the lack of protective histones, and higher coding density (Khrapko et al., 1997; Marcelino & Thilly, 1999). Recent research into mtDNA mutations has revealed that their frequency increases with age and varies among tissues (Sanchez-Contreras et al., 2023).

The retina, the multi-dynamic post-mitotic tissue responsible for vision, is comprised of many cell types, all heavily reliant on the functions of mitochondria for health and survival. Mitochondria have several essential roles that maintain cell health, including energy generation, regulation of calcium levels, and cholesterol and iron metabolism (Ferrington et al., 2020). Mitochondrial dysfunction in one or more retinal cell types has been implicated in human retinal degenerative diseases such as age-related macular degeneration (AMD), retinitis pigmentosa, diabetic retinopathy, and glaucoma (Eells, 2019). The retinal pigment epithelium (RPE) is a monolayer of cells located at the outer boundary of the retina and contains abundant mitochondria, perhaps due to its specific cellular functions. The RPE regulates the transport of nutrients from the choroidal blood supply to the neural retina and relies on reductive carboxylation and oxidative phosphorylation for its energy production (Du et al., 2016; Hurley, 2021). The RPE mitochondria become dysfunctional as we age, leading to an imbalance in this established symbiosis. This phenomenon is thought to be accelerated in disease states, with damaged RPE mitochondria driving the onset and progression of retinal degeneration or AMD (Kaarniranta et al., 2020). Investigation into the source of damaged RPE mitochondria has implicated the accumulation of mtDNA mutations as a source of mitochondrial dysfunction using normal and diseased human donor tissue. In addition, mtDNA mutation burden in the RPE appeared to be associated with both disease stage and likelihood of progression (Karunadharma et al., 2010). Cumulatively, these observations have provided a novel mechanism to explore in the context of retinal degenerative diseases.

The current study uses a previously developed mouse model of accelerated aging known as Polg^D257A^ mutator (PolgD257A) mouse to understand the role of mtDNA mutation accumulation in the retina. This model has a single amino acid substitution (D257A) in the exonuclease domain of the *Polymerase gamma* (Polg) protein responsible exclusively for mtDNA replication and repair. This mutation does not impact the replicative capabilities of Polg but impairs its ability to perform base-excision repair, resulting in an approximately 2500-fold increase in mtDNA mutations (Kujoth et al., 2005). Since its development, this model has been widely used as a tool to study aging and accelerated mtDNA mutation burden across various tissues, including brain, liver, and skeletal muscle (Dai et al., 2014; Hiona et al., 2010; Maclaine et al., 2020). However, the retina or RPE has never been investigated in the context of this mouse model. Mutations in Polg have commonly been associated with neurodegenerative diseases such as Alpers-Huttenlocher syndrome, childhood myocerebrohepatopathy, ataxia neuropathy spectrum, and autosomal progressive external ophthalmoplegia (Rahman & Copeland, 2019). Other diseases associated with specific mtDNA point mutations include but are not limited to Leber’s hereditary optic neuropathy, mitochondrial encephalopathy, and myoclonic epilepsy (Zeviani & Di Donato, 2004). By understanding how the error prone Polg protein and accumulation of mtDNA mutations impact the lifespan of a mouse, this study aims to elucidate their role in retinal aging. This study uses diverse and complementary methodologies to test the hypothesis that the accumulation of mtDNA mutations plays a causative role on the onset and progression of retinal degeneration. As the first study to employ the Polg^D257A^ mitochondrial mutator mouse with consideration for the retina, this analysis characterizes the effects of mtDNA mutations on mitochondrial structure, function, and retinal health.

## 2. RESULTS

### 2.1 Accelerated age-related morphological and functional retinal deficits in the PolgD257A mice

To investigate the role of mtDNA mutation burden on the retinal layers, optical coherence tomography (OCT) was performed using a previously established protocol (Singh et al., 2020). Imaging was performed on Polg^WT/WT^ (wild-type, WT), Polg^WT/D257A^ (heterozygous), and Polg^D257A/D257A^ (PolgD257A) littermate mice at 3-, 6-, and 9-months of age (Figure 1A). Raw images were then averaged, adjacent to, within 100um (??CHECK) of the optic nerve, and heat maps were generated and quantified using InVivo Vue Imaging Software (Figure 1B). Over the observed time course, the total retinal thickness of WT and heterozygous mice did not significantly change. However, the total retinal thickness of the PolgD257A mice was significantly decreased, with an average thickness difference of 10.27 µm (Figure 1C). Individual layers were quantified to identify which retinal cell layer were contributing to the observed decrease in retinal thickness. The outer nuclear layer (ONL), which contains the nuclei of the cone and rod photoreceptors, showed a significant decrease in thickness by 6 months of age and continued to be significantly decreased at 9 months of age (Figure 1D). The calculated average thickness difference was 2.96 µm at 6 months and 6.63 µm at 9 months accounting for ∼65% of the total decline in retinal thickness at both time points. Other retinal layers showing differences at specific ages included the RPE, photoreceptor outer segments (OS), and retina nerve fiber layer (RNFL) (Figure S1A). To determine if the mtDNA mutation burden had any effect on retinal cells’ response to light flashes, electroretinography (ERG) was performed using a previously established protocol (Aiello et al., 2023). PolgD257A mice showed significantly decreased a-wave amplitude at 6 months of age (Figures 1E and 1G), indicating a functional reduction in rod photoreceptor cells at this time. Additionally, PolgD257A mice showed decreased b-wave amplitude at both 6- and 9-months of age (Figures 1E and 1H), indicating a functional reduction in bipolar cells at these time points. Finally, the most drastic decrease in the PolgD257A mice’s retinal function was observed in the light-adapted response, significantly reduced at 6- and 9-month time points (Figures 1F and 1I). The light-adapted response is a physiological measure of cone photoreceptor cell health. As indicated by previous literature (Ball et al., 2022; Li et al., 2020), the cone photoreceptors rely significantly more on healthy mitochondrial function than their rod counterparts, giving context to the observed results in the PolgD257A ERG studies. Overall, these results indicate that PolgD257A mice experience retinal thinning at an earlier age than WT, which is most pronounced in the photoreceptor layer. Additionally, all recorded responses identify a functional compromise in the PolgD257A mice, with the most severe phenotype observed in the testing of cone photoreceptors. Since there were little to no observed significant differences between the WT and heterozygous compared to PolgD257A mice using both modalities, moving forward, PolgD257A mice were only compared to WT.

**Figure 1:**
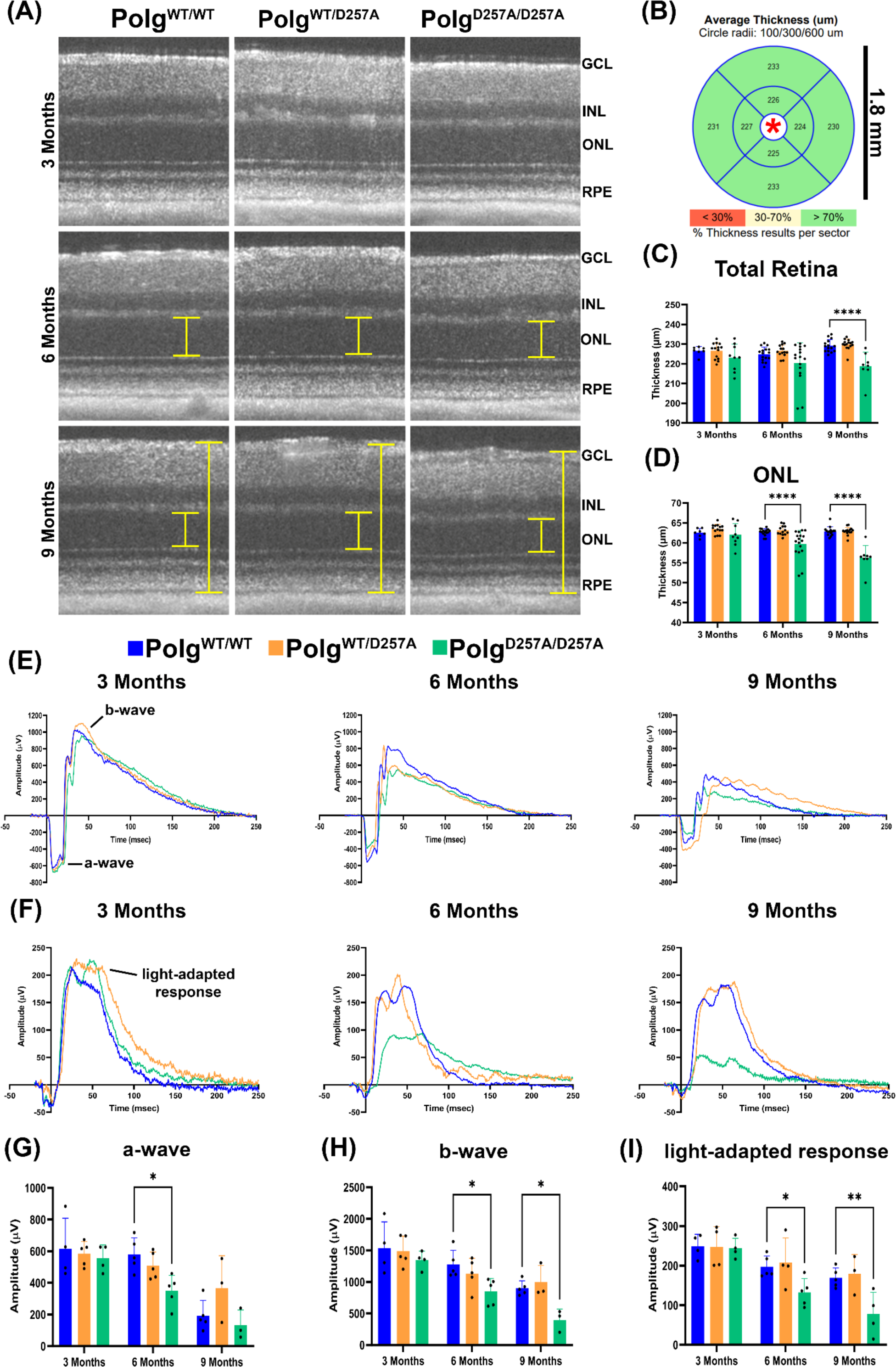
In vivo morphological and functional retinal characterization of PolgD257A retinas. (A) Averaged representative raw OCT image files (yellow bars = layers of retina where thickness differences were observed). (B) Representative heat map assembled by InVivoVue software used for downstream analysis (only green regions representing >70% thickness results per sector were quantified for accuracy) Quantification starts 100 µm from optic nerve head. (C) Graphical representation of total retinal thickness and (D) outer nuclear layer thickness (ONL). (E) and (F) Representative wave-form traces of dark-(E) and light-adapted (F) electroretinography (ERG) in response to a blue-green flash stimulus (1.4 log cd·s/m^2^) (G), (H), and (I) Amplitude of the a-wave (G), b-wave (H), and light-adapted response (I) in response to the flash luminance at 24.1 cd·s/m2. Data are expressed as mean ± SD. * p = 0.01 to 0.05; two-way ANOVA; Data points represent biological replicates; asterisks indicate significance.

### 2.2 Accelerated age-related changes in retina of the PolgD257A mice

Histology and immunohistochemistry with retinal cell markers were performed to gain a deeper understanding of the structural and molecular changes occurring in the PolgD257A retina. Mouse retinal histological sections stained with toluidine blue revealed a decline in total retinal thickness, particularly in the in the ONL of the photoreceptors by 9 months of age (Figures 2A, brackets, 2B, and 2C). Furthermore, analysis of PolgD257A mice retinas probed with an R/G opsin antibody, a commonly used marker for cone photoreceptor outer segments (Greenwald et al., 2014), showed significantly reduced labeling for this protein by 6 months of age. By 9 months the level of this protein was also reduced in WT mice, signifying that aging contributes to the loss of this protein over the mouse’s life (Figures 2D, arrows, and 2F). PolgD257A mice retinas were also probed with a rhodopsin antibody, a commonly used marker for photoreceptor outer segments (Park, 2014). A reduction in this protein was observed at 6 and 9 months of age, although neither was significant (Figures 2E, arrows, and 2G). To investigate retinal bipolar cells, an interneuron that creates a pathway from the photoreceptors to the ganglion cells, the protein kinase C alpha (PKCα) marker that stains both the inner and outer synaptic clefts was used (Ruether et al., 2010). The labeling for this protein also decreased with aging and decreased earlier and to a greater extent in the PolgD257A mice (Figures 2E and 2H). Interestingly, all markers showed increased expression at 3 months of age in the PolgD257A mice, possibly revealing a compensatory mechanism that is enhanced between the 3- and 6-month-old PolgD257A mice. This data validates previously observed in vivo results, again indicating a more severe phenotype in the cone photoreceptors.

**Figure 2:**
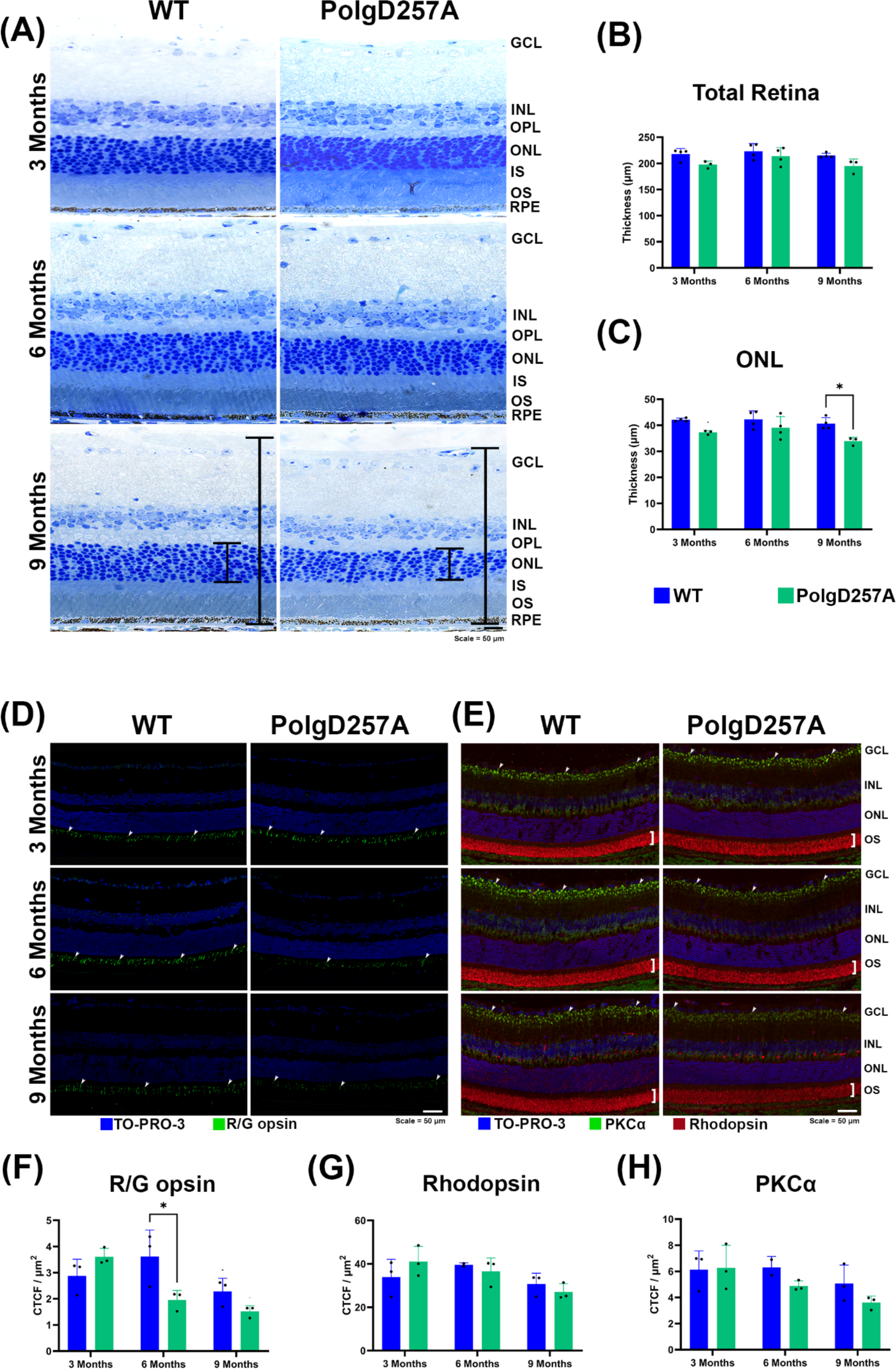
Loss of normal retinal morphology and essential proteins in the PolgD257A mouse. (A) Representative retinal histological sections stained with toluidine blue. (B) Graphical representation of measured total retinal thickness and (C) outer nuclear layer (ONL) thickness. (D) Immunofluorescence staining of cone photoreceptor outer segment marker R/G opsin. (E) Immunofluorescence staining of rod photoreceptor outer segment marker rhodopsin and bipolar cell synapse marker PKCα. (F) Graphical representation of R/G opsin staining, (G) rhodopsin staining, and (H) PKCα staining. CTCF = corrected total cell fluorescence. Data are expressed as mean ± SD. * p = 0.01 to 0.05; two-way ANOVA; Data points represent biological replicates; asterisks above for significance.

### 2.3 Accelerated age-related changes in RPE of the PolgD257A mice

During the previously described retinal immunohistochemistry studies, more autofluorescent granules were observed in the RPE cytoplasm. To understand this observation further, we compared higher magnification images to the relative amount of autofluorescence in the RPE cytoplasm. At 3 months, the PolgD257A showed an increased rate of autofluorescent granule accumulation in the RPE layer (Figures 3A, arrowheads, and 3B). By 6 months of age, the WT RPE displayed granule formation similar to the PolgD257A mice. However, PolgD257A there was a trend of increase in larger granules. By 9 months, this trend was significant, with PolgD257A RPE having significantly larger granules when comparing the top 10 granule sizes per mouse (Figures 3A and 3C). Differences in granule formation and accumulation became more apparent when comparing the summary statistics of WT to PolgD257A at each time point (Figure 3D). The average granule size increased with age and was more drastic in the PolgD257A mice at each time point. The most frequent granule size remained small (∼0.78 µm^2^). The median granule size and 75^th^ percentile also increased with age in both WT and PolgD257A mice. Interestingly, the maximum granule size was drastically increased in the PolgD257A mice, with at least one granule being 25.605 µm^2^ at 9 months of age (Figure 3D). To better understand the nature of the observed granules and to gain additional information on the RPE ultrastructure in the PolgD257A mice, electron microscopy analysis was performed at 3 months of age. The PolgD257A RPE showed disorganized microvilli (Figure 3E, MV) and disorganized basal infoldings (Figure 3E, BI). Additionally, PolgD257A pigment appeared smaller and disorganized when compared to the proper apical membrane alignment seen in WT (Figure 3E, P). Undigested photoreceptor outer segments near the basal surface were frequently observed in the PolgD257A RPE cytoplasm (Figure 3E, OS). Finally, PolgD257A RPE showed an increased frequency of electron-lucent vacuoles per cell (Figure 3E, white asterisk). This data suggests that the PolgD257A RPE accumulates more and larger autofluorescent granules with age, and changes in overall RPE morphology can be seen as early as 3 months.

**Figure 3:**
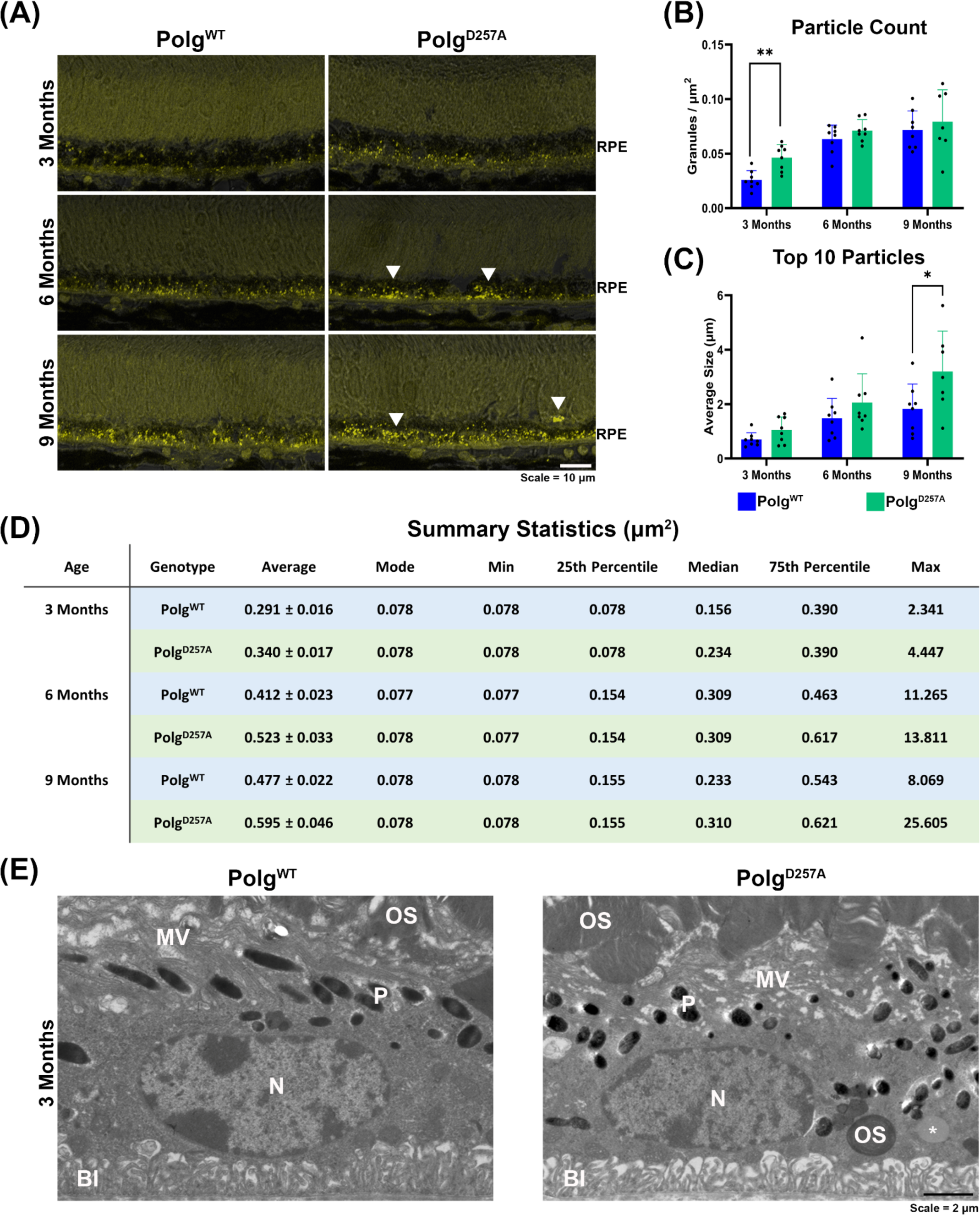
PolgD257A RPE exhibits altered auto-fluorescent granule accumulation size distribution with age. (A) Representative retinal sections excited between 488 – 594 nm wavelength with overlayed bright field to show RPE specific auto fluorescent granule accumulation. White arrowheads represent larger granules. (B) Graphical representation of the number of particles counted in the RPE layer after background thresholding divided by the area of RPE analyzed. (C) Graphical representation of the averaged top ten particle sizes per image acquired. (D) Summary statistic table of RPE auto fluorescent granule data (µm2). (E) Representative electron micrographs of 3-month-old WT and D257A mouse RPE (N = 3). MV = microvilli, P = pigment granule, N = nucleus, OS = outer segment, BI = basal infoldings. Data are expressed as mean ± SD. * p = 0.01 to 0.05; two-way ANOVA; Data points represent biological replicates; asterisks above for significance.

### 2.4 Higher levels of oxidative stress do not contribute to retinal accelerated age-related changes in the PolgD257A mice

Because many of the observations detected changes in the PolgD257A occurred at 3- and 6- months of age, we proceeded to gain a deeper understanding of the biochemical changes occurring in the retina and RPE of the PolgD257A at these ages, before aging became a compounding factor.

To start investigating the driving mechanisms behind the pathology of the PolgD257A retina and RPE, oxidative stress was measured through several methods. Previous studies on this mouse model have shown tissue-specific increases or decreases in compounds related to oxidative stress (Hiona et al., 2010; Zsurka et al., 2018). We performed a protein carbonylation assay to detect common protein oxidation products in biological samples, specifically carbonyl derivatives of proline, lysine, arginine, and threonine residues (Madian & Regnier, 2010). This assay measures 2, 4- Dinitrophenylhydrazine to identify protein carbonyl groups directly correlated to protein oxidation, as previously described (Bhattacharyya et al., 2022; Madian & Regnier, 2010). Retinal protein lysates of 3- and 6-month-old WT and PolgD257A mice were tested, with no significant increase observed in protein carbonylation (Figures 4A and 4B). Similarly, RPE lysates of 3- and 6-month-old mice, showed no significant difference in protein carbonylation between WT and PolgD257A (Figures 4C and 4D). Additionally, WT and PolgD257A retinal sections were probed for 8-hydroxy-2’ - deoxyguanosine (8-OHdG), a well-established biomarker for oxidative damage to both mitochondrial and nuclear DNA (Valavanidis et al., 2009). At 3 months of age, PolgD257A retinas showed a qualitative increase in this marker observed in the ganglion cell layer (GCL), consistent with previous literature describing the robustness of this staining in the mouse retina (Gu et al., 2022), although not significant (Figure 4E, arrows); minor 8-OHdG labeling is observed in the INL (Figure 4E, arrowheads). A qualitative increase in the GCL and INL labeling was observed in the 6-month-old WT retinas (Figure 4E, arrows, and arrowheads). The 6-month-old PolgD257A retinas showed a trend of decrease in 8-OHdG labeling in retinas compared to WT; however, the 8-OHdG labeling was qualitatively increased in the INL at this age (Figures 4E and 4F, arrowheads). Previous studies reported that the retinal 8-OHdG levels naturally increase with age (Wang et al., 2010), but this increase was not observed with aging in the PolgD257A mice We also analyzed the extent of 8-OHdG labeling in the RPE was analyzed. Again, no discernible differences were found between WT and PolgD257A mice. However, 8-OHdG did increase with age in both mice (Figures 4G and 4H). Overall, this data suggests that an increase in oxidative stress is not correlated with mtDNA mutation burden in the retina and RPE.

**Figure 4:**
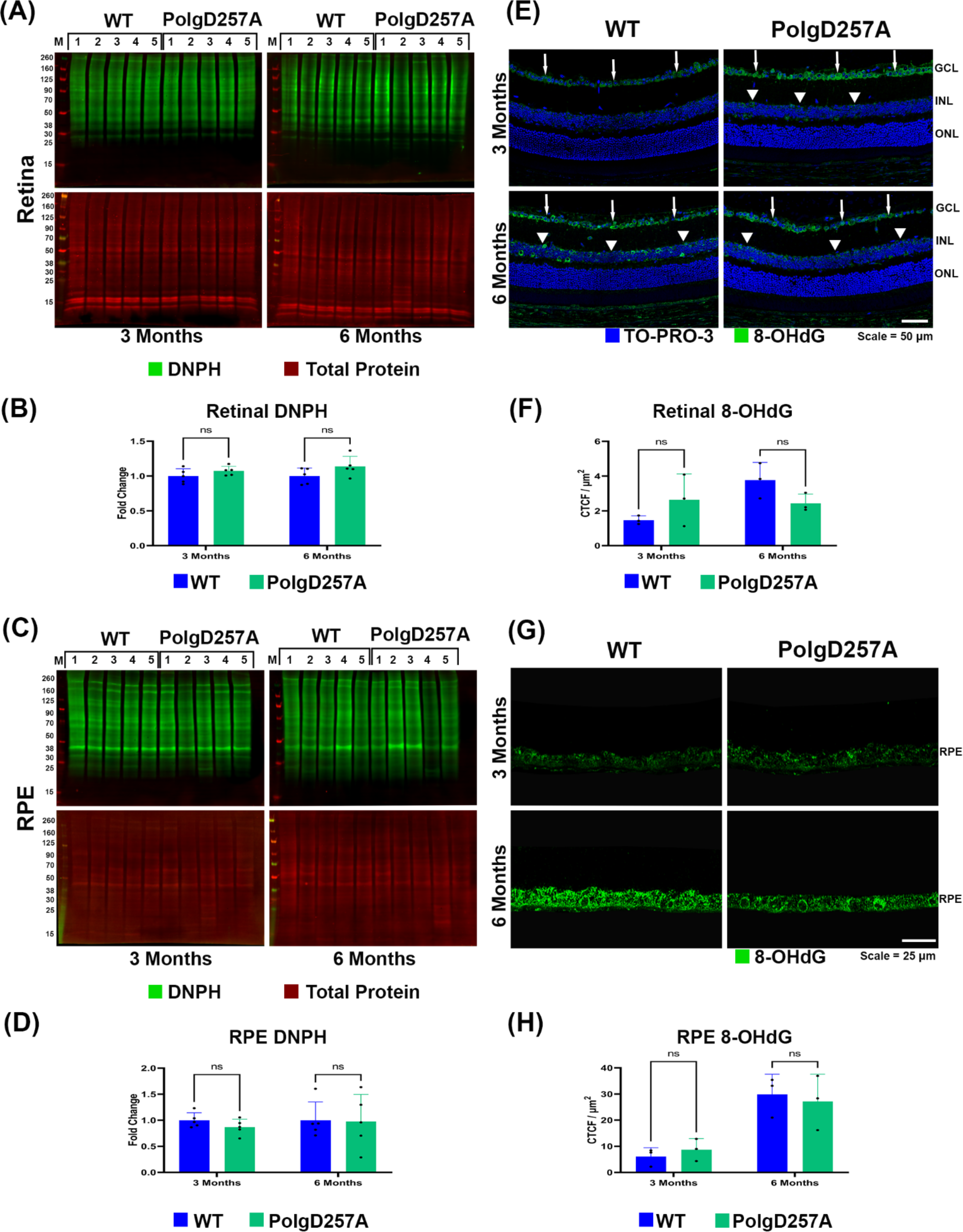
Oxidative stress is preferentially increased with age and not correlated to mtDNA mutation burden. (A) Protein carbonylation assay of WT and D257A total retina protein lysate. (B) Graphical representation of observed retinal DNPH (2, 4-Dinitrophenylhydrazine) normalized to total protein content. (C) Protein carbonylation assay of WT and D257A total RPE protein lysate. (D) Graphical representation of observed RPE DNPH normalized to total protein content. (E) Representative retinal sections stained with 8-OHdG (8-hydroxy 2’-deoxyguanosine). White arrowheads to show elevated staining in the GCL (ganglion cell layer) and INL (inner nuclear layer). (F) Graphical representation of retina 8-OHdG staining, CTCF = corrected total cell fluorescence. (G) Representative RPE sections stained with 8-OHdG. (H) Graphical representation of observed RPE 8-OHdG staining. Data are expressed as mean ± SD. * p = 0.01 to 0.05; two-way ANOVA; Data points represent biological replicates; asterisks above for significance.

### 2.5 Retinal and RPE mitochondrial dysfunction in the PolgD257A mice

Several mitochondrial-specific morphological and functional proteins were investigated to test the hypothesis that mitochondrial dysfunction was promoting the retinal phenotypes described above. First, western blot analysis of POLG protein was performed in retinal lysates to determine if the D257A mutation led to differential protein expression. POLG expression was not found to be changed between age or mtDNA mutation burden (Figures 5A and 5B). Voltage-dependent anion channel (VDAC) and pyruvate dehydrogenase (PDHA1) were investigated to look at specific mitochondrial markers. PDHA1 is a commonly used mitochondrial matrix marker and was found to be upregulated in 6-month-old PolgD257A retina (Figures 5C and 5D). The upregulation of PDHA1 is associated with the enhancement of the mitochondrial-mediated apoptosis pathway and inhibition of aerobic glycolysis, promoting oxidative phosphorylation (Sun et al., 2019). VDAC was found to be significantly upregulated at 3 months in the PolgD257A retina (Figures 5C and 5E); with an increasing trend at 6 months. In addition to being a commonly used mitochondrial outer membrane marker, this protein is also responsible for apoptosis by mediating the release of apoptotic factors from the mitochondria (Shoshan-Barmatz et al., 2020). To determine the overall consequence of PolgD257Ain photoreceptor ultrastructure, electron microscopy analysis was performed on WT and PolgD257A mice at 3 months of age. The PolgD257A cone inner segment (CIS) and rod inner segment (RIS) mitochondria appear more disorganized, less electron-dense, and improperly aligned with the cell body compared to WT (Figure 5F). A similar observation has been described when comparing old and young murine retina photoreceptors (Kam & Jeffery, 2015). Finally, to test the hypothesis that mtDNA mutation accumulation leads to dysfunctional components of oxidative phosphorylation complexes, NDUFB8 (complex I), SDHB (complex II), UQCRC2 (complex IIII), MTCO1 (complex IV), and ATP5A (complex V) were probed in retinal lysates by western blot (Figures 5G and 5H). The levels of complex V were not significantly changed at both ages in the PolgD257A retina (Figure 5I). Complex IV levels were significantly downregulated in the PolgD257A retina at 3 months of age compared to WT, but no significant changes were observed at 6 months (Figure 5J). The levels of complex III and complex II were not significantly changed in PolgD257A retinas (Figures 5K and 5L). Complex I levels were significantly reduced in the PolgD257A retinas at both ages investigated (Figure 5M).

**Figure 5:**
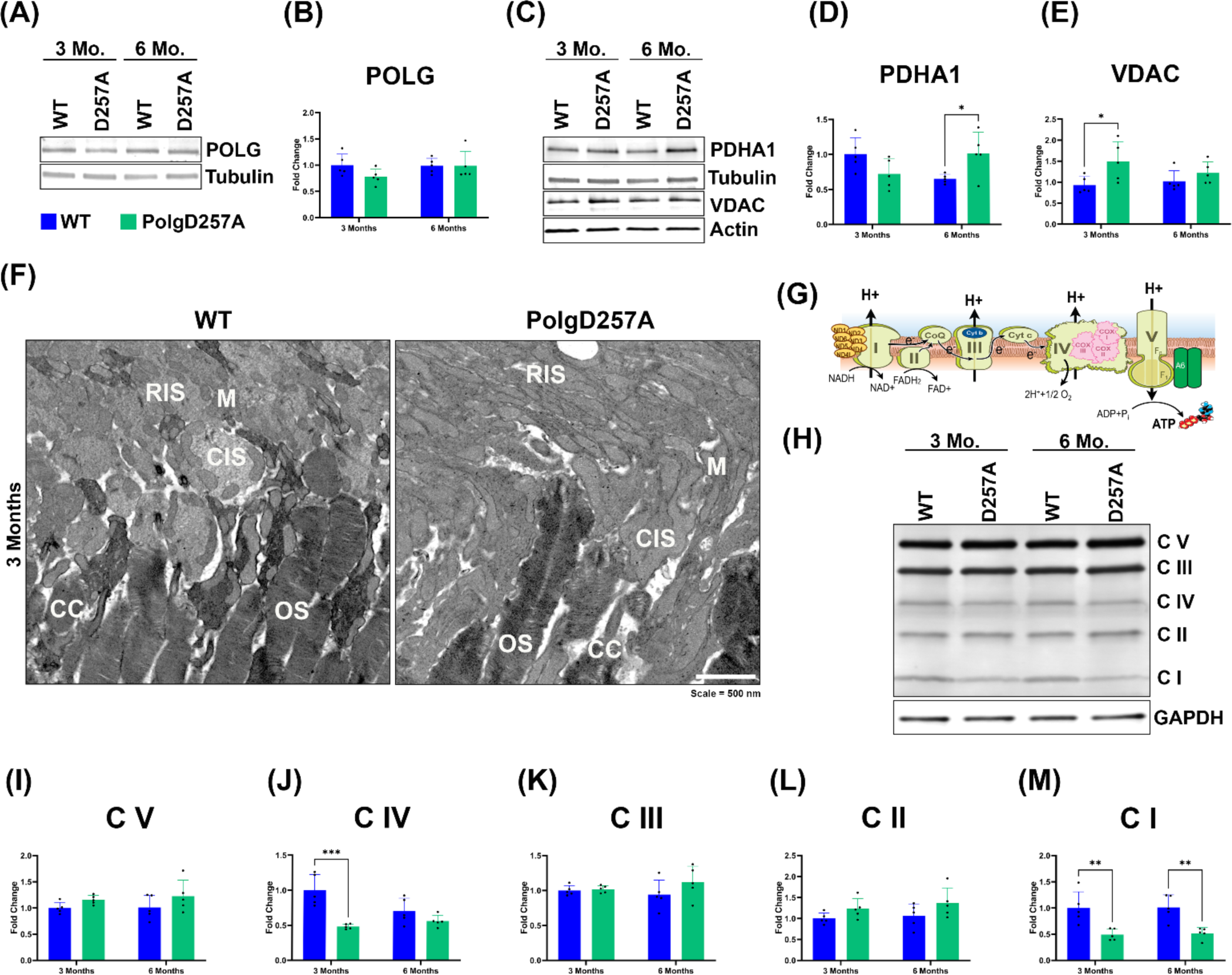
Loss of mitochondrial encoded complex proteins in the PolgD257A mouse retina. (A) Representative immunoblot of POLG protein content in the retina. (B) Graphical representation of POLG protein content in WT and D257A mice retina. (C) Representative immunoblot of PDHA1 and VDAC protein content. (D) Graphical representation of PDHA1 and (E) VDAC protein content. (F) Representative electron micrographs of cone photoreceptors in the WT and D257A inner segments. (G) Electron transport chain indicating mitochondrial encoded subunits of complex proteins. (H) Representative immunoblot of complex proteins (I-V). (I) Quantification of C V, (J) C IV, (K) C III, (L) C II, (M) C I normalized to GAPDH, and fold change calculated based off 3-month WT average. Data are expressed as mean ± SD * p = 0.01 to 0.05; two-way ANOVA; Data points represent biological replicates; asterisks above for significance.

The same experiments were performed using RPE protein lysate to determine if mitochondrial health was also being compromised in the RPE. Similar to the observed retina phenotype, POLG protein expression in the RPE was not significantly different between ages or in the PolgD257A mice (Figures 6A and 6B). Unlike the retinal phenotype, neither PDHA1 nor VDAC increased at either age in the PolgD257A mice RPE (Figures 6C, 6D, and 6E). Electron microscopy was performed to determine ultrastructural differences in RPE mitochondria. Mitochondria (M) in the PolgD257A mice appeared shorter, more rounded, and lacked densely packed cristae (Figure 5F). The basal infoldings (BI) appeared less dense and more disorganized in the PolgD257A RPE (Figure 5F). Finally, the PolgD257A RPE had a higher frequency of observable ribosomes in their cytoplasm (Figure 5F, arrowheads). Finally, components of oxidative phosphorylation complexes NDUFB8 (complex I), SDHB (complex II), UQCRC2 (complex IIII), COX1 (complex IV), and ATP5A (complex V) were probed in RPE lysates by western blot (Figures 6G and 6H). Quantification of complex V levels detected no changes between ages or genotypes (Figure 6I). Complex IV levels were significantly reduced in the PolgD257A RPE at 3 months and further reduced at 6 months. However, WT complex IV levels were significantly reduced with age (Figure 6J). There was decreasing trend for complex III and II in the 3-month PolgD257A RPE (Figure 6K). The levels of these complexes also displayed a trended decrease with age in the WT RPE (Figure 6L). Finally, complex I levels were significantly decreased in the PolgD257A 6-month RPE (Figure 6M). These data suggest that the observed changes in both retina and RPE are not due to increased or decreased POLG levels since expression of this protein remains stable with age and genotype. Additionally, the increased expression of PDHA1 and VDAC at specific ages in the PolgD257A retina could indicate apoptosis activation, explaining the significant loss in ONL thickness and cell count observed above. Both retina and RPE ultrastructural investigations indicate a loss of proper mitochondrial morphology and organization, as well as other indicators of declining overall cell health. Complex proteins I and IV are significantly reduced in the retina and RPE of the mutant mouse, suggesting a preferential loss of these proteins over others. These complexes contain the most mtDNA encoded subunits, with seven subunits in complex I and three in complex IV (Figures 5G and 6G, yellow/gray subunits). Since this mouse accumulates a rapid rate of mtDNA mutations, these subunits are likely preferentially disrupted, leading to an improper assembly of the entire complex. Finally, aging plays a significant role in the expression of complex proteins, specifically in the RPE. Most complexes (C IV, C III, CII) were significantly depleted between 3 months and 6 months in WT mice RPE, suggesting a higher susceptibility to aging-related oxidative phosphorylation complexes degeneration than the retina counterparts.

**Figure 6:**
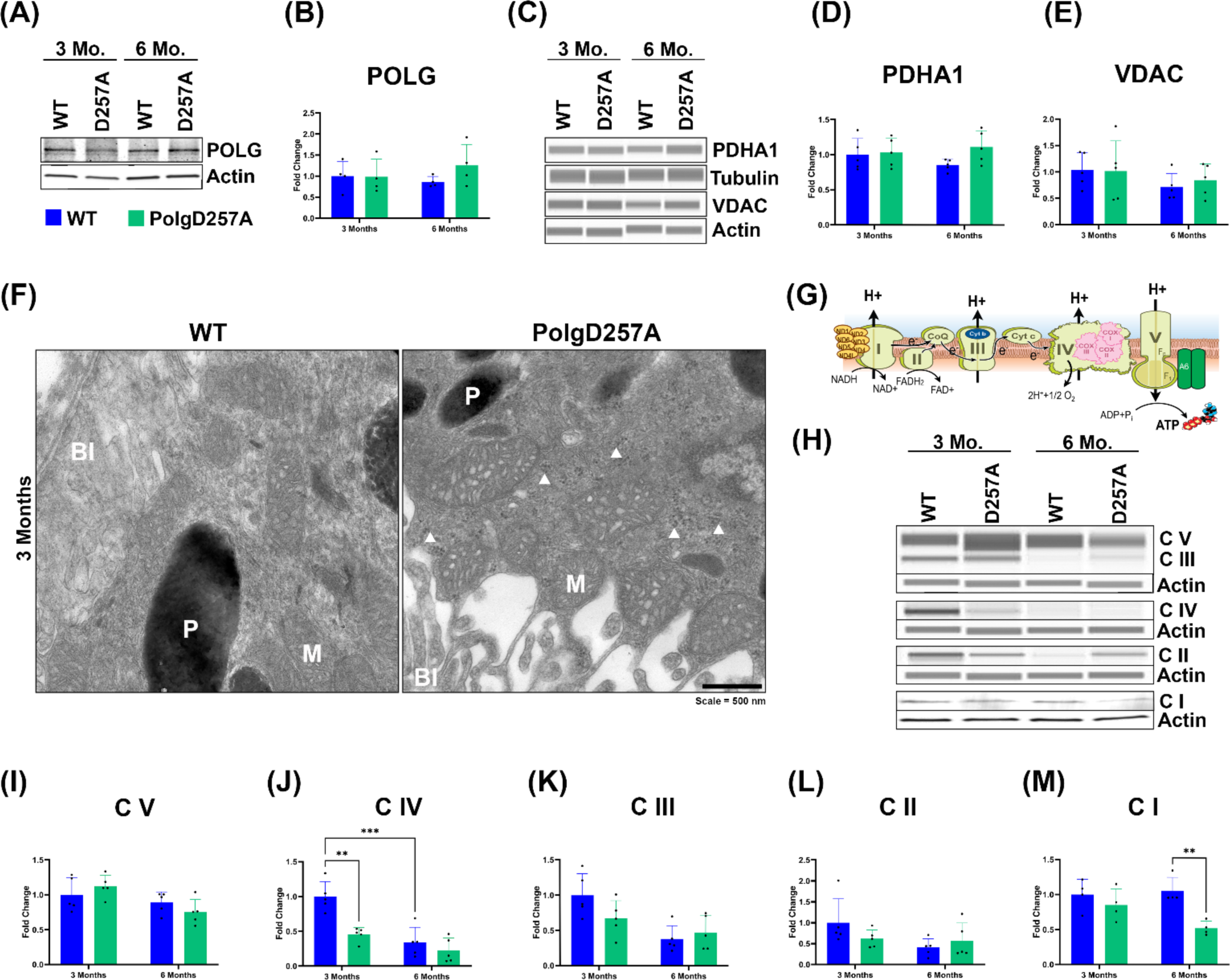
Loss of mitochondrial encoded complex proteins in the PolgD257A mouse RPE. (A) Representative immunoblot of POLG protein content in the RPE. (B) Graphical representation of POLG protein content in WT and D257A mice RPE. (C) Representative immunoblot of PDHA1 and VDAC protein content. (D) Graphical representation of PDHA1 and (E) VDAC protein content. (F) Representative electron micrographs of mitochondria in the WT and D257A RPE. M = mitochondria, P = pigment granules, BI = basal infoldings, R = ribosomes. (G) Electron transport chain indicating mitochondrial encoded subunits of complex proteins. (H) Representative immunoblot of complex proteins (I-V). (I) Quantification of C V, (J) C IV, (K) C III, (L) C II, (M) C I normalized to GAPDH, and fold change calculated based off 3-month WT average. Data are expressed as mean ± SD. * p = 0.01 to 0.05; two-way ANOVA; Data points represent biological replicates; asterisks above for significance.

## 3. DISCUSSION

The current study comprehensively examines the retina and RPE health of the PolgD257A mouse model. A wide range of ages was used to delineate the contributions of normal physiological aging processes with those attributed explicitly to mtDNA mutation accumulation. This is the first characterization in vivo and ex vivo of this model in the context of the retina and provides a working overview of baseline phenotypes. Our in vivo morphological and physiological retinal analysis detected little to no observed significant differences between the WT and heterozygous compared to PolgD257A mice. Our results are in agreement with a previous study that reported that despite harboring mtDNA mutation levels that were higher than those reached during normal aging in WT mice, heterozygous POLG mice were reported to have no “overt” phenotype (Vermulst et al., 2007).

We systematically demonstrate here that PolgD257A mice develop accelerated age-related progressive retinal abnormalities. A summary of the age-related changes identified is shown in Table 1. Our data indicate that PolgD257A retinas display signs of morphological abnormalities and physiological dysfunction associated with mitochondrial dysfunction. During aging, mtDNA repair pathways are limited and they begin to fail, leading to additional mtDNA damage and contributing to the development of retinal degenerative diseases such as glaucoma, diabetic retinopathy, retinitis pigmentosa, progressive external ophthalmoplegia, and age-related macular degeneration (Jarrett et al., 2008; Zeviani & Carelli, 2021). These results presented here establish the PolgD257A mice as a model to understand the contribution of mtDNA mutation burden in retinal degeneration.

**Table 1.**

| Methods | OCT | ERG | R/G Opsin | PKCα | Rhodopsin | RPE granules | TEM | Retina DNPH | RPE DNPH | Retina 8-OHdG | RPE 8-OHdG | Retina Complex proteins | RPE Complex proteins |
| --- | --- | --- | --- | --- | --- | --- | --- | --- | --- | --- | --- | --- | --- |
| 3 Months | No observable changes in total retinal thickness | No observable changes in cell function | Non-significant increase in staining | No observable changes in staining | Non-significant increase in staining | Significantly more granules / $\mu\text{m}^2$ | Disorganized microvilli, basal infoldings, and pigment; undigested OS and presence of vacuoles | No observable changes in protein carbonylation | No observable changes in protein carbonylation | Non-significant increase in staining | No observable changes in staining | Significant decrease in COX IV and COX I | Significant decrease in COX IV |
| 6 Months | Thickness decreases in ONL | Decreased a-wave, b-wave, and light adapted response amplitudes | Significantly decreased staining | Non-significant decrease in staining | Non-significant decrease in staining | Non-significant increase in granules / $\mu\text{m}^2$ and average granule size | N/A | No observable changes in protein carbonylation | Significantly more protein carbonylation with age but not genotype | Non-significant decrease in staining | Significantly more staining with age but not genotype | Significant decrease in COX I | Significant decrease in COX I |
| 9 Months | Thickness decreases in ONL and total retina | Highest observed decrease in b-wave and light adapted response amplitudes | WT loses staining, creating decreased staining difference | Non-significant decrease in staining | Non-significant decrease in staining | Significantly larger average granule size | N/A | N/A | N/A | N/A | N/A | N/A | N/A |

Our in vivo OCT analysis revealed total retinal thinning by 9 months of age, with significant thinning of the ONL of the photoreceptors observed at 6 and 9 months of age. This change, observed in vivo, could be validated by histology, and attributed to the ONL thinning identified in 9-month-old PolgD257A retinas.

The ERG testing detected significantly decreased a-wave amplitude at 6 months in PolgD257A mice, indicating a functional reduction in rod photoreceptor cells at this age. These results were corroborated by the reduced rhodopsin labeling observed in 6 and 9 months PolgD257A retinas since rhodopsin is the molecule that absorbs light and thus ‘senses’ light under dim light conditions (Park, 2014). The ERG b-wave amplitude was significantly decreased in 6 and 9 months PolgD257A retinas. The observed reduction of PKCα labeling corroborated these results in the PolgD257A mice as scotopic ERG b-waves reflect light-induced electrical activity in retinal cells post-synaptic to the photoreceptors and are primarily generated by the activity of depolarizing (ON) bipolar cells (Pardue & Peachey, 2014). In addition, the b-wave is also affected by OFF-center bipolar cells and by light-induced activity in amacrine and ganglion cells (Awatramani et al., 2001). The observed reduction of PKC in the PolgD257A mice suggest that bipolar cell health may be compromised. Finally, the cone ERG was significantly decreased at 6 and 9 months in PolgD257A mice. These results were corroborated by a significant decrease in the red/green cone opsin labeling suggesting a reduction in R/G cones may underlie the photopic ERG defect (Sondhi et al., 2021).

The investigation of the RPE layer of the PolgD257A mice showed a higher abundance of autofluorescent granule accumulation in their cytoplasm. However, these granules were not observed by TEM, suggesting that these granules are not lipofuscin. Lipofuscin is a long-lived intracellular inclusion body, lipid- and bisretinoids-rich, and autofluorescent material that progressively accumulates in the RPE during aging and pathological conditions (Feeney, 1978). Transmission electron microscopy micrographs of the lipofuscin granules exhibit uniform electron density and a size range consistent with previous results of lipofuscin granules in RPE sections (Ng et al., 2008; Schraermeyer & Heimann, 1999). Further experiments are needed to confirm the molecular nature of these autofluorescent granules. Our data suggests that the observed autofluorescent granules could be composed of something removed during tissue preparation for electron microscopy, such as retinoids (Kiser & Palczewski, 2016). The early changes observed in the RPE likely promote the unhealthy state of the retina at later time points. Our electron microscopy images detected additional changes, such as basal infoldings disorganization in the PolgD257A RPE. The changes in these RPE structures could be driving a nutrient deficit in the RPE and, therefore, a deficit in the neural retina (Bonilha, 2008). Because the RPE changes were observed before other changes in the retina, we hypothesize that the RPE dysfunction may promote the retinal dysfunctions. Previous studies on Polg257A mice reported alopecia, anemia, reduced subcutaneous fat, kyphosis, and osteoporosis by approximately 6 months (Trifunovic et al., 2004; Vermulst et al., 2007; Vermulst et al., 2008). Thus, the RPE dysfunction observed at 3 months of age suggests a crucial role of mtDNA mutations in these cells.

Interestingly, oxidative stress in the retina and RPE of this model does not appear to be elevated compared to WT but is elevated with normal physiological aging. Our data is in agreement with previous studies on young and aged PolgD257A mice, which did not detect a significant increase in levels of oxidative stress markers (Dai et al., 2014; Hiona et al., 2010; Kujoth et al., 2005; Mott et al., 2001; Trifunovic et al., 2005). Our data presents evidence against the “vicious cycle” of reactive oxygen species production and mtDNA mutagenesis (Hahn & Zuryn, 2019; Hiona & Leeuwenburgh, 2008), providing further proof that mtDNA mutations likely discourage the production of endogenous free radicals (Kennedy et al., 2013; Nissanka & Moraes, 2018). The accumulation of mtDNA mutations in the PolgD257A mouse model likely leads to a decrease in overall ETC complex activity, promoting a reduction in free radicals (Andreyev et al., 2005).

Finally, analysis of mitochondrial markers, morphology, and ETC complex proteins revealed substantial differences. The accumulation of PDHA1 and VDAC in the retina indicate the activation of the apoptotic pathway, signifying a method for the observed ONL thinning. The decline in cristae density and normal mitochondria shape in the RPE could also indicative declining mitochondrial function and energy production. Our data agrees with previously reported data from the PolgD257A mouse, supporting the hypothesis that mtDNA mutations can promote tissue dysfunction through the loss of critical irreplaceable cells due to activation of apoptosis (Kujoth et al., 2007).

Our western blot analysis revealed a preferential decline in complex proteins between PolgD257A and WT. Specifically, complex I and IV were most impacted in the retina and RPE of 3-month PolgD257A mice, while complex I was most impacted in the 6-month-old retina and RPE. Interestingly, these complexes contain the most mtDNA-encoded subunits, with seven and three proteins encoded by mtDNA. These data corroborate previously published data indicating that the primary impact of dysfunctional Polg encompasses a specific reduction in complexes I and IV (Hauser et al., 2014; McLaughlin et al., 2020). Previous studies also detected with a marked reduction in the content of ETC complexes in muscle cells (Hiona et al., 2010), intestine cells (Fox et al., 2012) and brain (Dai et al., 2013; Perier et al., 2013; Ross et al., 2019).

Although this study provides a comprehensive baseline characterization of many different retinal and RPE phenotypes in this mouse model, it fails to identify a specific functional mechanism behind the observed decline in retinal health. The most likely origin of the decaying retina is the implication of mitochondrial dysfunction, which is apparent in this mouse model. It is likely that mtDNA heteroplasmy, defined as the percentage of mutated mtDNA molecules compared to WT, plays a role in observed phenotypic variation (Parakatselaki & Ladoukakis, 2021). Since findings in this study do not support the long-standing “vicious cycle” theory of mitochondrial dysfunction and oxidative stress, the observed defects are possibly a consequence of impaired mitophagy, nutrient deprivation, or acute immune responses, which were not investigated. Mitophagy is the process by which dysfunctional mitochondria are expelled and replaced with normal functioning mitochondria. This process has already been thoroughly characterized as playing a role in RPE heterogeneity (Datta et al., 2023). The disruption of mitochondria observed at an early age in the RPE of this model could indicate a respiratory chain inhibition, and therefore, a nutrient deprivation in the neural retina that supports the idea of a “metabolic switch” where RPE cells start consuming metabolites initially intended to be supplied to the retina (Hurley, 2021; Zhang et al., 2021). Finally, damage to mtDNA in this model may lead to an acute immune response in the retinal tissue. The cGAS / STING pathway is a well-studied cytosolic DNA sensor system that has been identified to recognize mtDNA and initiate an acute immune response in these cells, leading to inflammatory-induced damage that can be therapeutically targeted (Kim et al., 2023; Zou et al., 2022).

Future studies of this model should include an investigation into all these well-studied mechanisms of retinal degeneration to determine the impact of mtDNA mutations in these pathways. An analysis of proteomic and metabolic differences in the PolgD257A mouse retina will provide further insight into the essential proteins and metabolic pathways that are disrupted in this mouse’s retina and RPE. Additionally, since there was a distinct preferential loss of complex proteins, mtDNA sequencing analysis of this tissue could provide more evidence for ETC dysfunction and how it is correlated to specific mtDNA point mutations in different regions of the mitochondrial genome. Finally, this model should be combined with other developed retinal degeneration models to identify its susceptibility to oxidative or light-induced damage (Koster et al., 2022; Wang et al., 2023). Overall, this study identifies the importance of the preservation of mtDNA in the aging mouse retina and the vulnerabilities that dysfunction of this system presents to retinal health.

## 4. METHODS

### 4.1 Mice

All procedures were approved by the Institutional Animal Care and Use Committee (ARC 2021-2191) of the Cleveland Clinic. Twelve-week-old PolgD257A and WT mice were purchased from the Jackson Laboratory (Bar Harbor, ME). Experiments were conducted on male and female littermate Polg^WT/WT^ (wild-type, WT), Polg^WT/D257A^ (heterozygous), and Polg^D257A/D257A^ (PolgD257A). Mice were genotyped as previously described (Kujoth et al., 2005). To not introduce mtDNA mutation burden during birth and development, male heterozygous Polg^D257A^ mice were crossed with C57Bl/6J females to generate wild-type (WT), heterozygous Polg^D257A/WT^ and homozygous Polg^D257A/D257A^ experimental mice. Mice were housed in individually ventilated cages in a 14-h light/10-h dark cycle and were provided regular chow and water ad libitum. Mice were tested, and tissue collected at ∼ 3 hours after light input to avoid circadian variations.

### 4.2 In vivo imaging of retinas

Imaging by spectral-domain optical coherence tomography (SD-OCT) (Leica Envisu R2210 UHR, Leica Microsystems) was carried out following sodium pentobarbital anesthesia and pupil dilation with 1.5 μL of 0.5% Mydrin-P drops (Santen Pharmaceutical Co., Ltd., Osaka, Japan) and topical anesthesia 0.5% proparacaine HCl as previously described (Singh et al., 2020) High-resolution volumetric SD-OCT imaging was used for age-based monitoring of retinal layer thickness. Linear b-scans (1.8 mm, 1000 a-scans × 20 frames × 1 b-scan) and volumetric rectangular scans (1.8 mm × 1.8 mm, 500 a-scans × 500 b-scan × 3 frames) were captured using 55° mouse OCT lens. Thickness measurements of each eye were performed using automated mouse retina segmentation software (Diver 1.4, Bioptigen Inc., InVivoVue Imaging Software). The scan parameter used for heatmaps were 1.8 mm diameter scan, 500 b-scans, 500 a-scans/b-scans, 3 frames average. The volume intensity projections (VIP) is divided into grids (sectors around the optic nerve head) and 9 x 9 grid locations are used for thickness measurements. The retina VIP is then segmented into eight classical retinal layers and thicknesses of each layer are measured by the InVivoVue software at 9 x 9 (i.e., 80 locations, ignoring the 81st location at the centra of the ONH). The software then averages the points falling in one sector of ETDRS grid and generate the heat map report. The heat map report contains - averaged heat map, segmented volume intensity projections (VIP) of the layers and average thickness ETDRS grid of the layers. Grid consists of three concentric circles of radii 100µm, 300 µm, and 600 µm. Average thickness is calculated for segmented points contained in the sector. Results for innermost sector (circle of 100 µm radius) are not calculated because all scans were centered on the optic nerve head. The resulting heat maps (thickness map) were analyzed for all eight retinal layers visible in OCT, which included the retinal nerve fiber layer (RNFL), inner nuclear layer (INL), inner plexiform layer (IPL), outer plexiform layer (OPL), outer nuclear layer (ONL), inner segment (IS), outer segment (OS), and RPE as previously reported (Singh et al., 2020). These measurements were performed in eight sectors divided around the optic nerve.

### 4.3 In vivo testing of retinal function

Retinal functions of mice were assayed after overnight dark adaptation as previously described (Aiello et al., 2023). Mice were anesthetized with 65 mg/kg sodium pentobarbital. Eye drops were used to anesthetize the cornea (1% proparacaine HCl) and to dilate the pupil (2.5% phenylephrine HCl, 1% tropicamide, and 1% cyclopentolate HCl). Mice were placed on a temperature-regulated heating pad throughout the recording session. In brief, electroretinograms (ERGs) were recorded in response to strobe flash stimuli presented in the dark by an Espion E3 ColorDome Full Field Ganzfeld (Diagnosys, Lowell, MA). An Ag/AgCl electrode in contact with the cornea was referenced to a needle electrode placed in the mouth of the mouse, and a ground lead was placed in the tail. Scotopic responses were obtained in the dark with ten steps of a blue-green flash stimulus, ranging from −3.6 log cd.s/m^2^ to 2.1 log cd.s/m^2^. The duration of the inter-stimulus intervals increased from 4 seconds for low-luminance flashes to 90 seconds for the highest stimuli. Two minutes following the scotopic ERG, a 10-second blue-green stimulus (5 cd/m^2^) was presented to elicit the c-wave. After 7 minutes of light adaptation, cone ERGs were recorded with strobe-flash stimuli (−1 to 2 log cd·s/m^2^) superimposed on the adapting field. Amplitude of the a-wave was measured at 8.3 milliseconds following the stimulus. The b-wave amplitude was calculated by summing the amplitude of the a-wave at 8.3 milliseconds with the peak of the waveform after the oscillatory potentials (≥40 milliseconds). Light-adapted response amplitudes were calculated by summing the peak of the waveform with the amplitude at 8.3 milliseconds.

### 4.4 Histology and transmission electron microscopy (TEM) of retinas

Enucleated eyes were fixed overnight at 4 °C by immersion in 2% paraformaldehyde, 2.5% glutaraldehyde, and 5% CaCl2 in 0.1 M cacodylate buffer. After removing the anterior segments under a dissecting microscope, eyecups were processed for epon embedding as previously described (Bonilha et al., 2015). For bright-field microscopy, semi-thin sections were cut using a diamond histotech knife (DiATOME, Hatfield, PA), collected on glass slides, and stained with toluidine blue. Sections were imaged with a THUNDER 3D Assay inverted microscope equipped with a Leica K3C Color camera (Leica Microsystems, Germany). High-magnification images were acquired within 200 μm of the optic nerve head (on both sides). Images were exported to ImageJ software (NIH, Bethesda, MD) and calibrated using an embedded reference scale. The retinal layer thickness was delineated as delineated using the freehand line and measured in triplicate to obtain a mean thickness for each mouse. Three measurements were performed throughout each acquired image within 2 sections/eye. For TEM, the same block of epon-embedded samples ultra-thin sections of 85 nm were cut with a diamond knife, stained with uranyl acetate and lead citrate, and observed with a Tecnai G2 SpiritBT, electron microscope operated at 60 kV.

### 4.5 Retinal immunohistochemistry

Enucleated eyes were fixed overnight at 4 °C by immersion in 4% paraformaldehyde in D-PBS and sequentially infused with sucrose and Tissue-Tek O.C.T Compound (4583; Sakura Finetek, Tokyo, Japan). Cryosections (8 μm) were cut on a cryostat HM 505E (Microm, Walldorf, Germany) equipped with a CryoJane Tape-Transfer system (Leica Inc., Buffalo Grove, IL). For labeling, sections were washed with PBS, blocked in PBS supplemented with 1% BSA (PBS/BSA) and 0.1% Triton- X100 for 30 min, and incubated with primary followed by secondary antibodies coupled to Alexa fluorophores. Nuclei were labeled with TO-PRO-3 (Molecular Probes, Inc., Eugene, OR). Sections were imaged using a laser scanning confocal microscope (Leica TCS-SP8, Exton, PA) using the same acquisition parameters for each channel in the Leica confocal software LAS-X. Antibodies used included: PKCα (Santa Cruz, sc-8393, 1:200), Rhodopsin (Abcam, ab5417, 1:1000), Red/Green Opsin (Sigma, AB5405, 1:500), 8-OHdG (Santa Cruz, sc66036, 1:100), MTCO1 (Life Technologies, 459600, 1:250) and secondary antibodies coupled to Alexa 488 or 594 (Invitrogen, A11034, A11029, A11037). Autofluorescence of unlabeled cryosections was performed and analyzed using confocal microscopy in the green channel (FITC filter: 490 nm excitation/519 nm emission) and red channel (TRITC filter: 550 nm excitation/570 nm emission). Autofluorescence was overlaid on differential interference contrast (DIC) images. Signal intensity was quantified within 200 μm of the optic nerve head, and the corrected total cell fluorescence (CTCF) was calculated for each area according to the following formula: CTCF = integrated density – (area of selected cell x mean fluorescence of background readings).

### 4.6 Protein extraction and western blotting

Mechanically isolated retinas and RPE were lysed in RIPA buffer (Alfa Aesar, Havermill, MA) containing a protease and phosphatase inhibitor cocktail (Sigma-Aldrich). Retinas were sonicated twice for 15 s while RPE was passed through a 27 1/2 G syringe needle, incubated on ice, and vortexed every 5 min for 20 min. Both lysates were centrifuged for 10 min at 14000 rpm at 4 °C, after which the supernatants were collected for western blotting. Protein quantification was performed using the MicroBCA Kit (ThermoFisher Scientific), followed by retinal protein (30 μg) separation via 4–20% Novex Tricine SDS- PAGE (ThermoFisher Scientific) and transfer to PVDF membranes (Immobilon- FL; Merck Millipore; Burlington). Membranes were incubated with: Total OXPHOS Rodent WB Antibody Cocktail (ab110413, 1:250), VDAC Polyclonal Antibody (Invitrogen, PA1-954A, 1:1000), PDHA1 Monoclonal Antibody (Invitrogen, 9H9AF5, 1:1000), DNA Polymerase gamma Rabbit mAb (AbClonal, A1323, 1:500) followed by washing and incubation with anti-mouse IRDye®680RD, anti-rabbit IRDye®680RD, and anti-mouse IRDye®800CW (all from LI-COR Biosciences, Lincoln, NE). Immunoreactive signals were visualized using Oddessey CLx (LI-COR Biosciences). The protein levels were quantified using ImageJ software. β- actin (Cell Signaling, 8H10D10 1:5000), Tubulin (Cell Signaling, 2144, 1:5000), or GAPDH (Proteintech, 60004-1-Ig, 1:5000) was used as internal control and 3-month-old Polg^WT^ mice averages were used to calculate fold changes. For RPE samples that do not contain high protein content, Jess Simple Western was used to conserve protein lysate. Jess, an automated western nano-assay system (ProteinSimple, Bio-techne, CA, USA) was used with internal standards to quantify the expression of different proteins in RPE lysates. The lysates were diluted with 0.1x sample buffer and mixed with fluorescent standards and 400 mM of dithiothreitol to reach a final concentration of 0.9 µg/µl. 2.7 µg of total protein was loaded from each sample onto the Jess assay plate. The 12-230 KDa fluoroscence separation module was used in this study with the 25-well capillary system. Antibody diluent, anti-mouse NIR and anti-rabbit NIR from ProteinSimple were used to set up the assay plate, as recommended by the instrument manual. Primary antibodies corresponding to beta-actin (Cell Signaling, 8H10D10), VDAC (Invitrogen, PA1-954A), PDHA1 (Invitrogen, 9H9AF5), Total Oxphos (Abcam, ab110413) were used at 1: 10, 1: 25, 1: 10 and 1: 5 dilutions respectively. The samples were separated at 375 V for 25 min and subjected to 30 min of blocking, 90 min of primary antibody incubation and 60 min of secondary antibody incubation. Compass for simple western v.6.3 software was used to analyze the data.

### 4.7 Protein carbonylation detection

Mechanically isolated retinas and RPE were lysed in RIPA buffer as described above. Protein carbonyl residues in were quantified in 10 μg of the cell lysates using the Protein Carbonyl Assay Kit (Abcam, Boston, MA, USA, ab178020). Derivatization and neutralization were carried out according to the manufacturer’s protocol. A parallel set of lysates were treated with the 1x Derivatization Control solution. Individual oxidized proteins were resolved by SDS-PAGE on 4-20% polyacrylamide gel gradient gel (BioRad, 5671093). Resolved proteins were transferred to PVDF membrane at 75 V for 2h. The membranes were blocked using 5% skimmed milk in PBS for 2 h and incubated with primary anti-DNP rabbit antibody (1:5000) overnight at 4 °C, followed by incubation with anti-rabbit IRDye® 800CW. The same membranes were next incubated with REVERT total protein stain (LI-COR, 926-11011). Signal intensity in the derivatized lanes and the control lanes was normalized with total protein signal in the respective lanes.

### 4.7 Statistical Analysis

Data were analyzed using GraphPad Prism v10.1.1 (GraphPad Software, La Jolla, CA) and are presented as the mean ± standard deviation (SD). Two-way ANOVA with Tukey’s Multiple Comparisons test and unpaired, two-tailed Student’s t-test were used to determine statistical significance between groups with an alpha value of 0.05. P values ≤ 0.05 were considered statistically significant.

## ACKNOWLEDGMENTS

The authors thank Benjamin A. Routhier and Caroline Milliner for their technical assistance and Mei Yin for her assistance with the ultra-thin sections and electron microscopy. The authors also thank David Schumick, BS, CMI for preparation of the illustration included in the manuscript.

---

**Supplemental Figure S1A:**
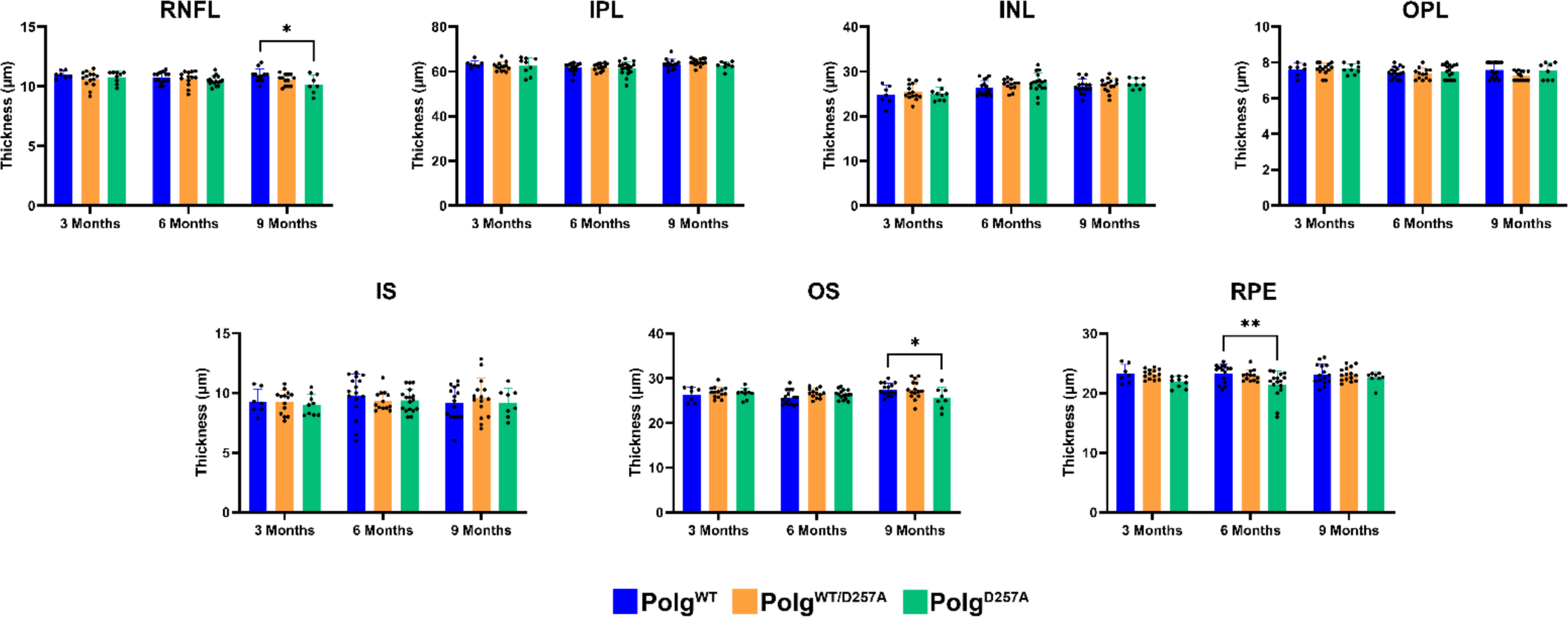
Retina layer thickness differences in PolgD257A determined by OCT. RNFL = retina nerve fiber layer; IPL = inner plexiform layer; INL = inner nuclear layer; OPL = outer plexiform layer; IS = inner segments; OS = outer segments; RPE = retinal pigment epithelium. Data are expressed as mean ± SD. * p = 0.01 to 0.05; two-way ANOVA; Data points represent biological replicates; asterisks above for significance.

