## Supplementary figures and images for "Modeling aging and retinal degeneration with mitochondrial DNA mutation burden"

### Supplemental Figure 1

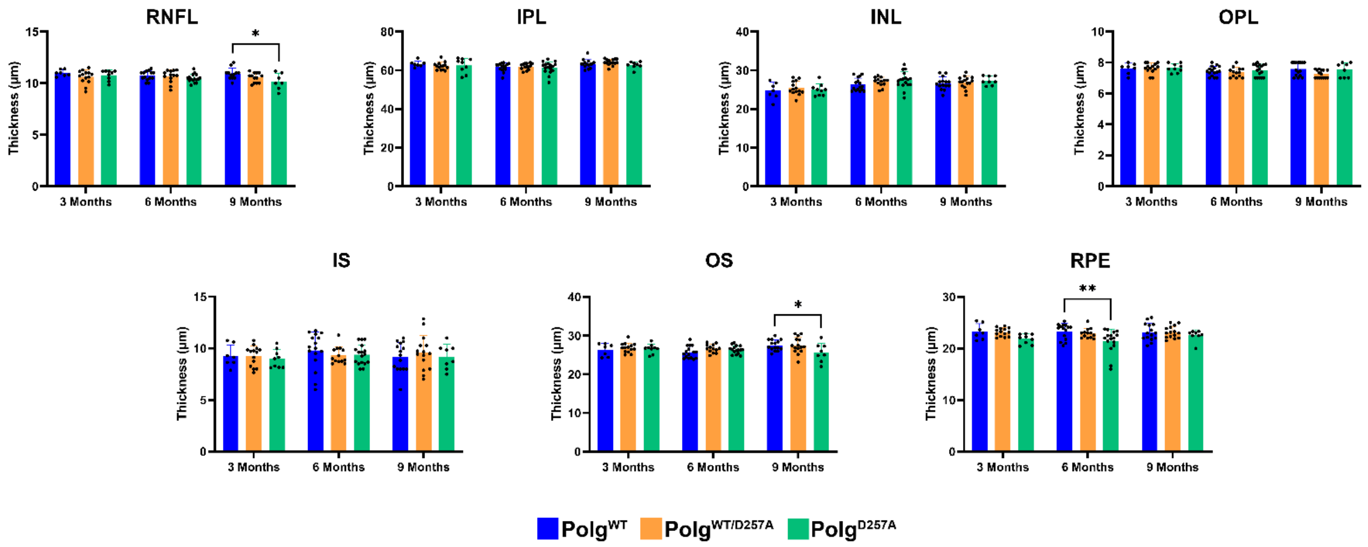
